# Two wild *Spinacia* species, *S. turkestanica* and *S. tetrandra*, genomes reveal a sex chromosome turnover in the genus

**DOI:** 10.1101/2023.11.08.566342

**Authors:** Hongbing She, Zhiyuan Liu, Zhaosheng Xu, Helong Zhang, Jian Wu, Xiaowu Wang, Feng Cheng, Deborah Charlesworth, Wei Qian

**Author notes:** These authors contributed equally to this work (Hongbing She, Zhiyuan Liu). **Correspondence:** Wei Qian, Deborah Charlesworth; Feng Cheng.

## Abstract

Spinach (*Spinacia oleracea*) is a dioecious species (with male and female flowers on separate individuals). Spinach and its closest wild relative, *S. turkestanica*, has homomorphic sex chromosomes, but the more distant relative *S. tetrandra* has heteromorphic sex chromosomes. We report high-quality genome assemblies for *S. turkestanica* and *S. tetrandra*. These diverged approximately 6.3 million years ago (Mya), while *S. turkestanica* split from *S. oleracea* much more recently, around 0.8 Mya, supporting previous suggestions that *S. turkestanica* is the direct progenitor of cultivated spinach. Using a combination of genomic approaches, we identified a sex-linked region (SLR) of ∼133 Mb in *S. tetrandra*. In all three species, the SLRs are within a large pericentromeric region of chromosome 4. We describe evidence that, in *S. tetrandra*, this region has completely stopped recombining in male meiosis, creating a large Y-linked region (YLR) that has partially degenerated; loss of recombination appears to have evolved in two events that created two “evolutionary strata”, one of which and is highly rearranged, relative to the X. The SLRs of *S. turkestanica* and *S. oleracea* are much smaller: both include only a 10 Mb Y-specific region which is not detected in *S. tetrandra*. This was duplicated into a 14 Mb inverted region, and is termed the Y-duplicated region, or “YDR”. These findings suggest that a turnover event created the YDR before these species diverged, replacing an extensive ancestral Y-linked region like the *S. tetrandra* YLR.

## Main text

The genus *Spinacia* is well suited for studying sex chromosome evolution. Cultivated spinach and its two close wild relatives, *S. turkestanica* Iljin and *S. tetrandra* Stev. (Ribera et al., 2020) are all dioecious. Although they are all diploids with 2n =12 (Fujito et al., 2015), and all have genetic sex-determining systems, and male heterogamety, their sex chromosomes differ. Phylogenic inferences using nuclear and chloroplast genome sequences demonstrated that *S. turkestanica* is closely related to *S. oleracea*, and both have homomorphic sex chromosomes (She et al., 2022a; Xu et al., 2017). The more distantly related *S. tetrandra* has heteromorphic sex chromosomes, with the Y larger than the X, based on flow cytometry results (Fujito et al., 2015).

In cultivated spinach, the YY genotype is viable (Wadlington and Ming, 2018), suggesting that the Y-linked region is physically small and not genetically degenerated, and/or evolved too recently for genetic degeneration to have occurred. The expected small size was recently confirmed when the fine structure of the X- and Y-linked regions in cultivated spinach (which we term XLR and YLR, respectively, or SLR collectively) were described (Ma et al., 2022; She et al., 2023). The YLR is 10 Mb larger than the XLR, due to presence of a Y-specific region carrying duplicated sequences of autosomal progenitors (termed the “Y duplication region”, or YDR) within a 14.1 Mb region in which genes are shared with the X, but which is arranged in inverted order in the X and Y assemblies (Li et al., 2023; She et al., 2023). Short-read resequencing data from a set of cultivated spinach individuals, plus both wild spinach species, showed that the *S. turkestanica* SLR is very similar to that of *S. oleracea*, including sharing similar TE insertions within the YDR (She et al., 2022b).

Genome regions carrying sex-determining genes have repeatedly evolved suppressed recombination. An extensive non-recombining region may thus have evolved in a common ancestor of all three *Spinacia* species, creating an ancestral heteromorphic XY chromosome 4 pair like that in *S. tetrandra* that subsequently underwent a turnover, as outlined in possibility (i) scenario (ii) includes several alternative possibilities. A small ancestral recombinationally inactive YLR, like that in spinach, might have expanded in the *S. tetrandra* lineage after the split from the other *Spinacia* lineage. For instance, expansion could occur by accumulation of repetitive sequences, though such a process seems unlikely to create a very large Y-linked region, which is necessary to account for the heteromorphism observed in *S. tetrandra*. A large sex-linked region is more likely to reflect a process in which more genes become included in the Y-linked region; this is predicted to occur if a sexually antagonistic polymorphism becomes established at a site closely linked to an ancestral SLR, creating selection for closer linkage to the SLR. This will be followed by divergence of the sequences of the new fully Y-linked genes, and can create new “evolutionary strata” like those observed in humans and other mammals (Bellott and Page, 2021; Lahn and Page, 1999; Skaletsky et al., 2003). Within such strata, repetitive sequences are predicted to accumulate, and genetic degeneration may eventually occur. *Spinacia* species may therefore shed light on the evolution of completely non-recombining sex chromosome regions.

To understand which of these scenarios has acted in *Spinacia*, we describe new chromosome-scale genome assemblies of *S. turkestanica*, and of the more distant spinach outgroup, *S. tetrandra*, which turns out to have a very large Y-linked region. They all have high transposable element (TE) densities, especially long terminal repeat elements (LTRs), which form 56.21% and 59.54% of the two wild spinach genomes (Table 1, left-hand column), similar to previous estimates for *S. oleracea* (She et al., 2023; Xu et al., 2017).

**Table 1.** TE densities within different genome region of the three *Spinacia* species, expressed as percentages of the regions indicated that are made up of TE sequences). The different regions of the sex chromosomes, and their locations in the different species, are explained below. Separate assemblies, allowing separate estimates for the XLR and YLR in *S. turkestanica*, are not available, and therefore the X size is not known, but, as the sex chromosomes are not heteromorphic, its size is very similar to the Y size.

| Species | Genome region |  |  |  |  |
| --- | --- | --- | --- | --- | --- |
|  | Whole genome | PARs | SLR | XLR | YLR |
|  |  | Sex chromosome regions |  |  |  |
| *S. tetrandra* | 75.92% | 70.78% | 77.49% | 75.78% | 80.84% |
| *S. turkestanica* | 74.36% | 74.44% | 85.05% | Not estimated separately |  |
| *S. oleracea* | 73.93% | 74.41% | 86.24% | 82.59% | 86.24% |

*S. turkestanica* is the closest wild relative of *S. oleracea*, and these species share a similar SLR on chromosome 4 (She et al., 2022b). We used synteny analysis to describe the *S. turkestanica* SLR in detail. Again very like *S. oleracea*, this species has a 24.8 Mb region that includes a 10.1 Mb YDR flanked by regions that are inverted in *S. oleracea* but within which *S. turkestanica* sequences align well (Table 1, Supplementary Table 1, Fig. 1). For the more distantly related species, *S. tetrandra*, we identified the sex-linked region *de novo*, independently of those of *S. turkestanica* and *S. oleracea*, using multiple approaches. First, normalized read coverage, estimated using seven sequenced males and five females, found lower values in males than females across a roughly 133 Mb region of chromosome 4, or about 61% of its assembly size (Fig. 1a, b, e, f). This result shows that chromosome 4 shows sex-linkage in this spinach species also, and that males are the heterogametic sex, as in the other two species. However, unexpectedly, the sex-differentiated SLR region in *S. tetrandra* is much larger than the SLRs of either *S. oleracea* or *S. turkestanica*, which, as explained above, are similar to one another (Table 1, Supplementary Table 1). Moreover, the low coverage in males suggests that the *S. tetrandra* Y chromosome includes an extensive partially degenerated region (with coverage in males low, but not as low as half that in females, see further details below), consistent with the species’ observed sex chromosome heteromorphism (Fujito et al., 2015). The pseudo-autosomal regions (PARs) total only 85 Mb, which, although large, is less than half the sizes in the other two species (Table 2). PAR2, at the left of this SLR, occupies about 63.0 Mb, or 29% of the chromosome, and the PAR near the right-hand end of the chromosome, PAR1, is about 22.1 Mb (10% of the chromosome, Fig. 1, Supplementary Table 1). *F_ST_* analysis confirmed the large sex-differentiated region inferred from coverage differences between *S. tetrandra* males and females. Using 82,053 SNPs in sequences that are detected in both the male and female individuals (i.e. not hemizygous in males), *F_ST_* analysis detects genetic differentiation (SNP frequency differences) between the sexes across about 133 Mb of chromosome 4 (Fig. 1c, g). Many SNPs in this region are homozygous in females and heterozygous in males (Fig. 1h, i), confirming the male heterogamety in *S. tetrandra* that is suggested by the coverage values.

**Figure 1.**
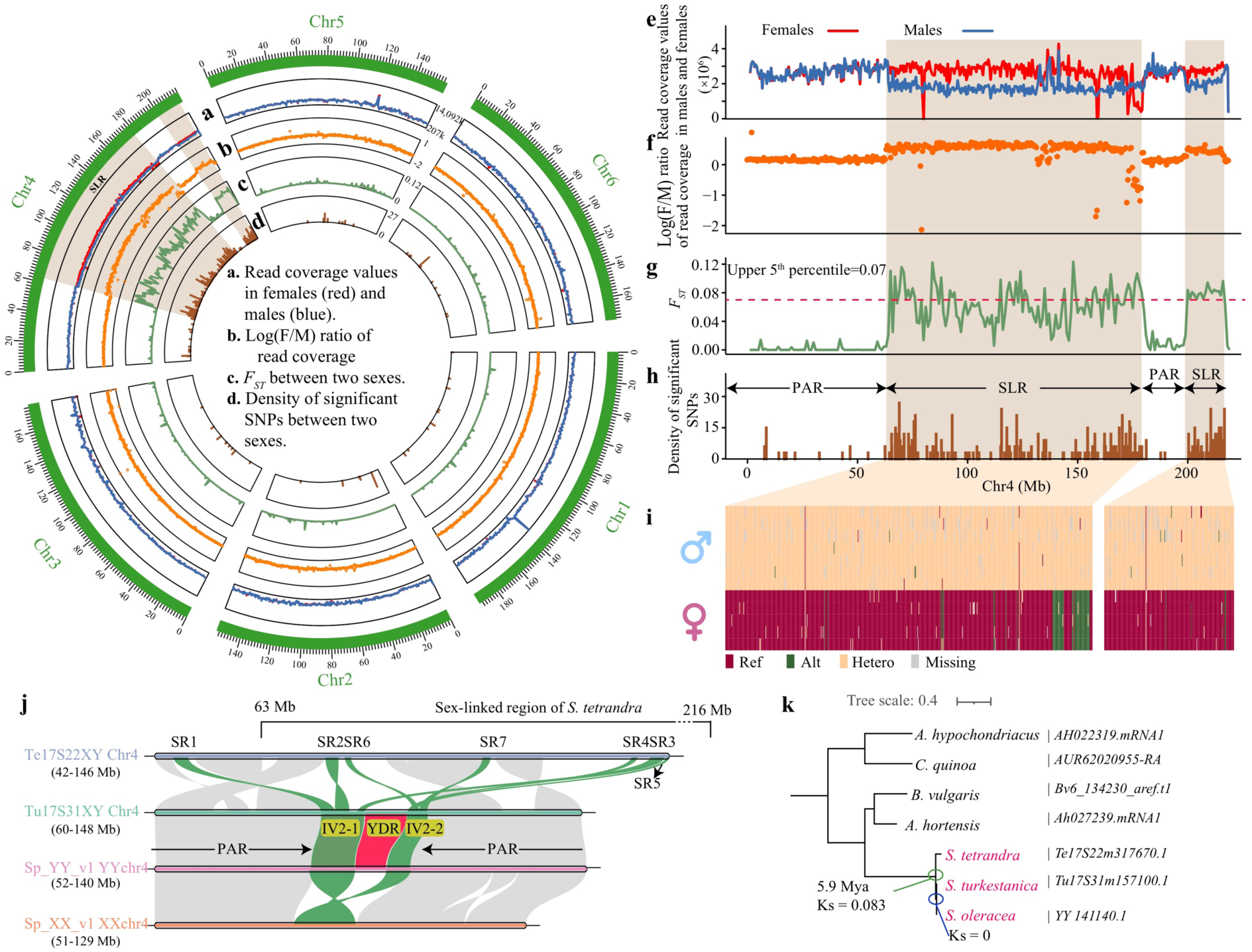
Identification of Chr4 as the sex chromosome, and of sex-linked region (SLR) in the *S. tetrandra* (Te17S22XY) assembly. **(a)** Read coverage values based on means of 5 females (red) and 7 males (blue) in 500-kb windows. **(b)** Ratios of read coverage, shown as Log (F/M) values. (c) *F_ST_*values between females and males. **(d)** Density of SNPs with allele frequency differences in the two sexes (in 1-Mb windows, using *P* < 0.01 in Fisher’s exact tests). The light brown background indicates the two regions described in the text that have the properties expected for SLRs, and the white regions are inferred to be pseudo-autosomal. **(e–h)** Expanded view of chromosome 4, showing the quantities corresponding to **(a–d)** in the Circos plot, using the same window size. **(i)** Haplotypes within the SLR inferred using 743 SNPs that differ between the sexes. Ref: reference allele; Alt: alternative allele; Hetero: heterozygous. **(j)** Synteny analysis of the SLR in cultivated spinach and its two wild relatives, using MCscanX. **(k)** Phylogenetic tree of the 414 bp *YY_141140.1* gene and its orthologs in the two wild spinach species and several outgroup species.

**Table 2.**
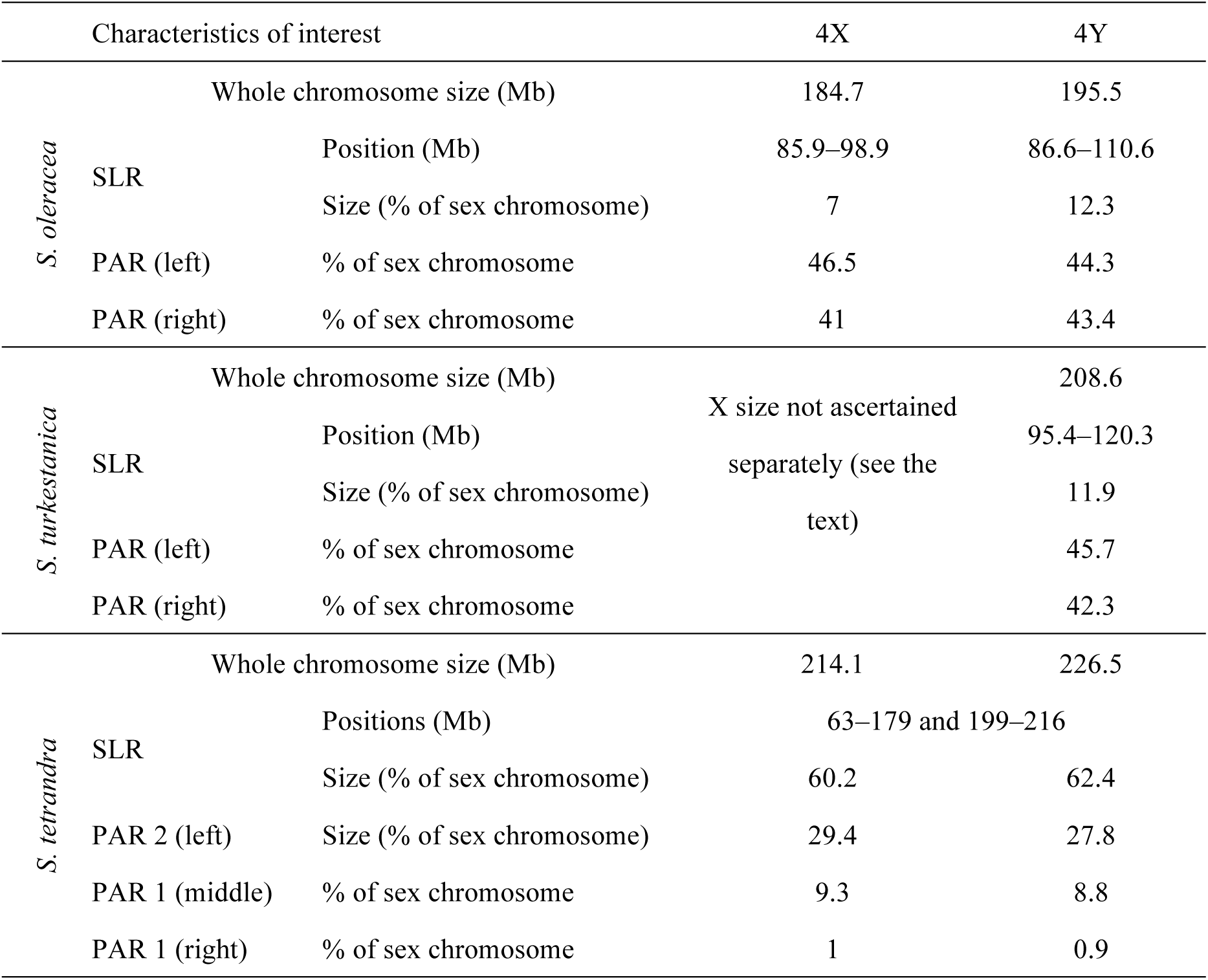
Characteristics of the sex-linked and pseudoautosomal regions in the three *Spinacia* species. The whole chromosome sizes are for the entire chromosomes, and include the different sized X- and Y-linked regions, denoted by SLRs in the table, and the PARs. In the text, we also refer to the long arm of chromosome 4, the sex chromosomes, as 4q. Note that, as explained above, the *S. turkestanica* X was not assembled separately but its size is similar to the Y size.

We identified seven syntenic regions shared between the SLRs of all three *Spinacia* species, the female *S. oleracea* assembly Sp_XX_v1, the *S. turkestanica* (Tu17S31XY assembly and the *S. tetrandra* assembly. All of these correspond to parts of the small (roughly 14 Mb) inverted regions described above in the regions flanking the YDRs of *S. oleracea* and *S. turkestanica*. In *S. tetrandra,* they are found in widely different locations scattered across the chromosome 4 assembly, probably reflecting rearrangements during the time when this lineage was evolving independently of the other *Spinacia* species (Fig. 1j, Supplementary Tables 2 and 3); rearrangements are discussed further below. We found orthologs of 35 *S. oleracea* YDR genes in the *S. tetrandra* assembly, but they are scattered across the genome (Supplementary Table 4), consistent with the previous conclusion that most *S. oleracea* YDR genes were recently translocated into the region (She et al., 2023). We previously described complete male-specificity of the *YY_141140.1* gene (coding sequence length: 414 bp) within the YDRs of both *S. oleracea* and *S. turkestanica* (She et al., 2022b). A phylogenetic tree of *YY_141140.1* and its orthologs shows that the relationships of the four outgroup species do not correspond with the species tree (Fig. 1k, Supplementary Table 5). Sequence divergence of the *YY_141140.1* and its homologous in the two species suggest that the genes of *S. turkestanica*/*S. oleracea* originated from *S. tetrandra* approximately 5.93 million years ago (Mya), based on molecular clock rate of 7.0 × 10^-9^ (She et al., 2022a; Xu et al., 2017). This is shortly after the split from *S. tetrandra* (estimated above to be 6.3 Mya).

In *S. oleracea* and *S. turkestanica*, the PARs occupy all of chromosome 4 other than the 24 Mb SLRs. We previously showed that the *S. oleracea* SLR is within a large pericentromeric region with high repetitive sequence densities, occupying about 127.1 Mb in the middle region of the metacentric sex chromosome. Similar repeat-rich regions are found at one end of each acrocentric autosome, probably indicating their centromere positions (She et al., 2023). Genetic mapping in *S. oleracea* showed that recombination is rare in these regions (She et al., 2023), and this can account for their high repetitive sequence densities in both the autosomes and the XY pair. Most of the PARs of this species are therefore recombinationally inactive, or recombine rarely, at most. In Figure 2, the terminal recombining regions (PAR 1 and PAR 2) are denoted by rPAR, and the pericentromeric regions that recombine less frequently are denoted by pPAR.

**Figure 2.**
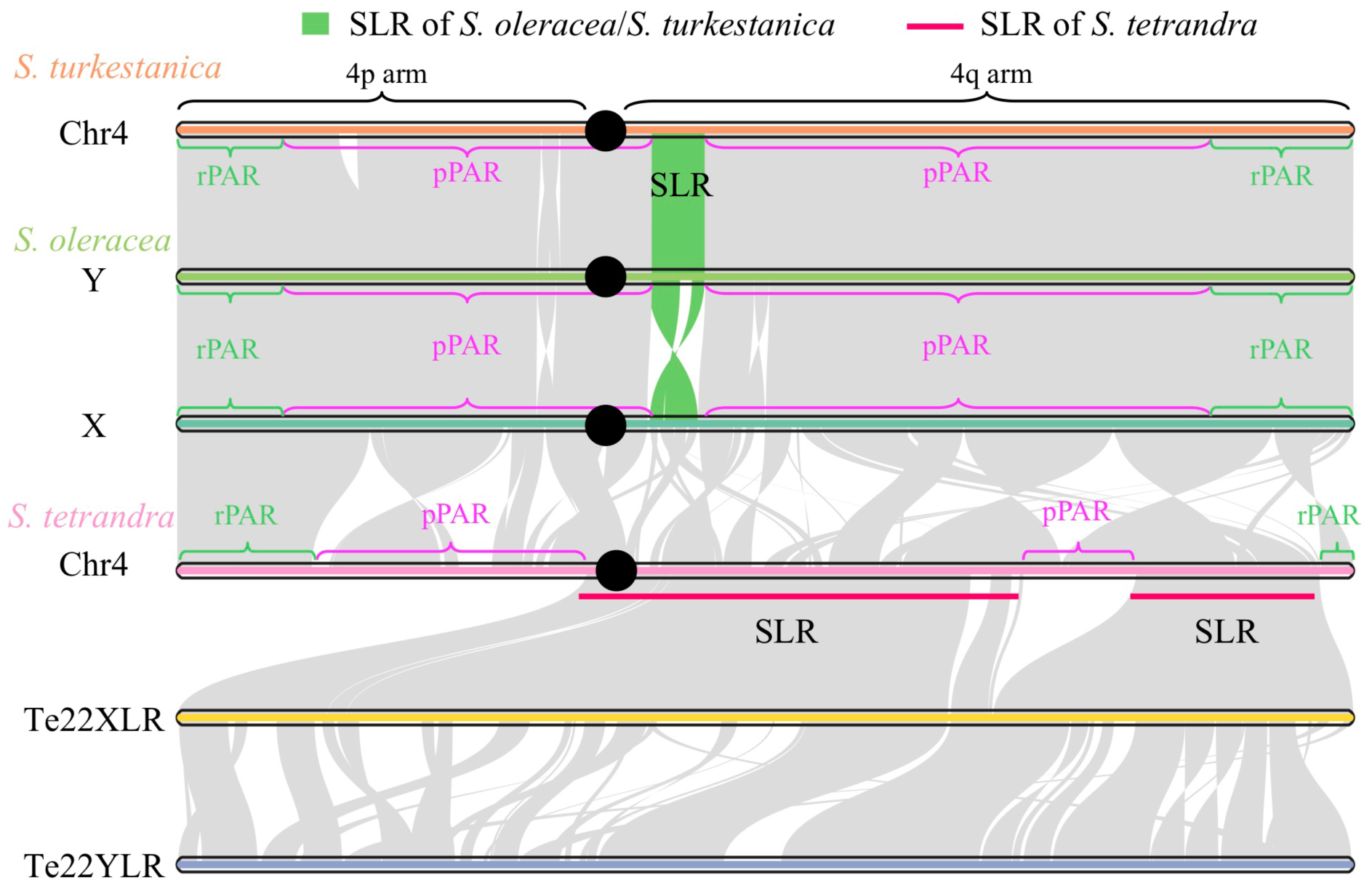
Comparison of sex chromosomes and among the three *Spinacia* species. Synteny analysis of the sex chromosomes of the three *Spinacia* species. The X and Y chromosomes of *S. oleracea* were from Sp_XX_v1 and Sp_YY_v1 assemblies, respectively (She et al., 2023). The black dot represent centromere position (see Method). SLR, sex-linked region; XLR, X-linked region, YLR, Y-linked region; PAR, pseudoautosomal region; rPAR, recombining PAR; pPAR, pericentromeric PAR. The rPAR and pPAR of *S. oleracea* were identified based on genetic mapping (She et al., 2023).

The two wild spinach relatives also have extensive gene-poor/TE-rich regions in the middle of the sex chromosomes, but the coverage and *F_ST_* results just described show that their sex-linked regions differ greatly. In *S. turkestanica* the SLR is a small region within the extensive pericentromeric region (Table 1, Fig. 2), as in *S. oleracea* (She et al., 2023), and chromosome 4 of this species can also be divided into regions corresponding to the spinach pPAR and rPAR; these regions are almost collinear in these two species. The same general gene density pattern is evident in *S. tetrandra*, but much of the region corresponding to the pericentromeric pPARs of the other two species is completely Y-linked and has undergone genetic degeneration, with many X-linked genes being hemizygous in males; the arrangement of the region is also very different from that in the other two species (Fig. 2). Due to the larger Y-linked region in *S. tetrandra*, this species’ PARs occupy only about 85 Mb in total, mostly at the two ends of the chromosome 4 X or Y assembly (Table 1, Fig. 2).

The shorter chromosome 4 arm (the left-hand arm) appears to be similar in all three species. Its PAR in *S. tetrandra*, here termed PAR2, is collinear with the left-hand rPAR of the other two species, and the size of this recombinationally active region is similar in all the species; it is also collinear in the *S. tetrandra* X and Y. The rest of PAR2 corresponds to part of the other species’ pPAR (this part is rearranged in *S. tetrandra*, consistent with this region being a rarely recombining region in this species also). The longer right-hand arm of the *S. oleracea* chromosome 4, which we term “4q”, however, appears to be different in *S. tetrandra*. Its PAR, termed PAR1, again corresponds to large portions of the other species’ 4q pPARs, but it also includes a part of their rPARs. Both of these regions are rearranged between *S. tetrandra* and the other two species (**Fig. 2**), reflecting the time during which these species have been evolving independently. However, the *S. tetrandra* 4q rPAR is much smaller than in the other two species. This suggests that, during the evolution of the *S. tetrandra* XY pair, this region is not descended from a pericentromeric region that rarely recombines in either sex. Instead, suppressed recombination in males evolved across a formerly recombining terminal chromosome region.

Genome-wide transposable element (TE) densities (mostly LTRs) are similar across all three spinach species, ranging between 73.9% and 75.9% (Table 1, Supplementary Table 6). In *S. turkestanica* and *S. oleracea*, the large PARs include extensive pericentromeric regions that rarely recombine (the pPARs), as well as regions at the ends of the two chromosome arms that recombine more often, albeit still rarely (the recombining PARs, or rPARs); in these PARs, TE densities are similar to the genome-wide values, because the large pericentromeric regions present on all the chromosomes dominate the estimates (Supplementary Table 6). The much higher TE density in the *S. turkestanica* and *S. oleracea* SLRs (above 85%, see Table 1, Supplementary Table 6) is consistent with TE accumulation after recombination became suppressed in the *S. turkestanica*/*S. oleracea* lineage at the time when the duplication that created the YDR.

The level of differentiation between two sexes in *S. tetrandra* allowed us to phase X and Y haplotypes by using male-specific SNPs (Supplementary Fig. 2). This yielded an X haplotype of 129.0 Mb and a larger (141.4 Mb) Y one, consistent with the observed heteromorphism (Supplementary Table 7). The phased X- and Y-linked regions (termed Te22XLR and Te22YLR, respectively) were further confirmed by Hi-C interaction, read coverage in the two sexes (Supplementary Figs. 3 and 4). The phased sequences allow us to test whether the YLR has accumulated TEs to a higher level than that of the XLR. The loss of genes from the YLR indicates that most or all of the chromosome 4 pericentromeric region is completely recombinationally inactive, so that repetitive sequences are expected to accumulate in the YLR after recombination stopped with the X. As predicted, the *S. tetrandra* YLR has a higher TE density than the XLR (Table 1, Supplementary Fig. 5, Table 8), as was also observed in the small *S. oleracea* YLR (She et al., 2023). The *S. tetrandra* YLR consequently has a lower gene density (16/Mb) than the XLR (24/Mb) (Supplementary Fig. 6, Table 7), or the other pericentromeric regions, which also have low gene densities. This repetitive sequence accumulation can explain the larger size of the Y than the X haplotype (**Fig. 3**). However, as described in the next section, the Y expansion is only seen in part of the SLR. In *S. tetrandra*, two LTR types of TEs, Copia and Gypsy, plus LTR elements classified as unknown types, make up most of the repetitive sequences in these species’ SLRs (Supplementary Table 6). Copia and Gypsy show similar densities across most of the X- and Y-linked regions, and similar overall mean densities in both the X and Y, but the unknown LTR types have consistently higher YLR than XLR densities (Supplementary Fig. 5, Table 8).

**Figure 3.**
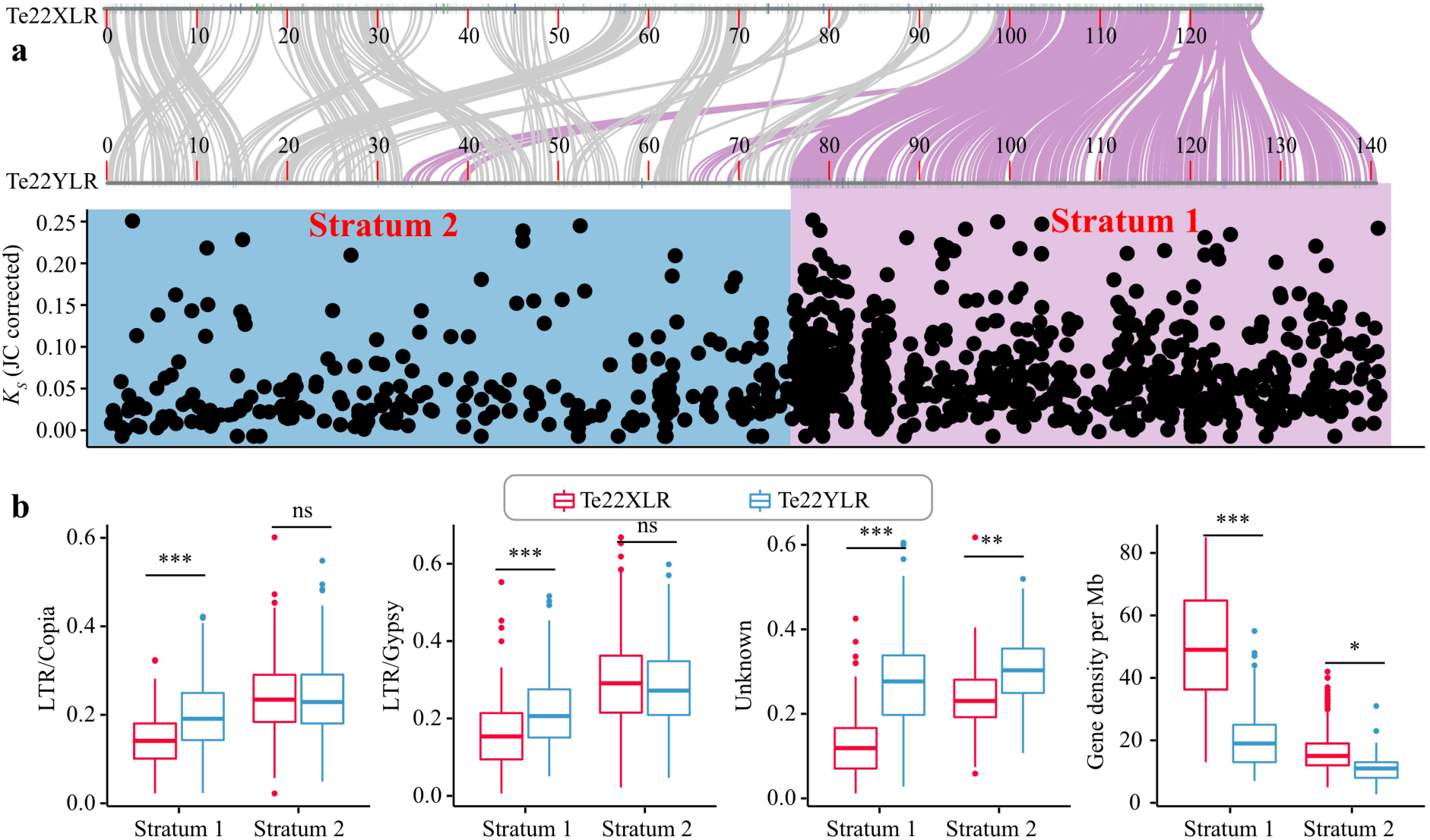
Evolutionary strata of the *S. tetrandra* sex chromosomes. **(a)** Synteny and *K_S_*values of allele pairs of genes found in the Te22XLR and Te22YLR sequences. The coordinates of the sequences are shown on diagrams of the XLR and YLR (and therefore both start at 0; the sizes of the PARs to their left and right of these regions are shown in Table 2 and Figure 1). Stratum 1 (pink), with the higher *K_S_*values and therefore probably the greater age, is considerably larger in the YLR than in the XLR, whereas for the inferred younger stratum 2 (blue) the size of the XLR is larger than that of the YLR. Small numbers of genes with higher *K_S_* values are not shown, as these sequences are saturated and their values are not reliable (Supplementary Fig. 8) The purple lines indicate the genes within stratum 1 on the XLR and their counterparts in YLR. **(b)** Boxplots showing the densities of three types of LTR elements in 200-kb windows, and gene density in 1 Mb windows, estimated within the two inferred strata of both XLR and YLR. The horizontal lines show the median values, and the boxes span the region from the 25 to the 75% quartile values. The whiskers extend to 1.5× the interquartile range. \*\*\**P*<0.001, \*\**P* <0.01, \**P*<0.05, ns, no significance.

If recombination is suppressed at different times, this forms evolutionary strata. To determine whether evolutionary strata are present in *S. tetrandra*, we examined sequence divergence between genes present on both the XLR and YLR. We used these data to estimate synonymous site divergence (*K_S_* values) for 1,525 homologous genes in the *S. tetrandra* XLR and YLR (Supplementary Table 9). The overall median *K_S_*value for the SLR genes is 6.8% (with Jukes-Cantor correction for saturation), much higher than in *S. oleracea* (median *K_S_* =1.6% for genes flanking the YDR; Supplementary Fig. 7; She et al., 2023). Change-point analysis of the *K_S_* values revealed two evolutionary strata within the SLR. (Fig. 3a, Supplementary Table 10). Stratum 2 includes 99.7 Mb at the left-hand end of the XLR (adjoining the region defined above as the PAR 2 end of chromosome 4; its YLR counterpart is 76.1 Mb). Its median Y-X *K_S_* value is 0.045. The rest of the XLR forms stratum 1, whose median *K_S_* value is 0.074, almost twice as high, and whose YLR is much larger than its XLR (65.2 Mb, about 46.1% of the 141 Mb completely Y-linked region detected in this species) (Fig. 3a, b). The regions occupied by these two putative strata in *S. tetrandra* differ greatly in their recombination rates, and such differences correlate with properties, including gene density, GC content, and the extent of DNA methylation (Rodgers-Melnick et al., 2015). The stratum that we infer to be younger, Stratum 2, corresponds entirely to regions that are pericentromeric or centromeric in the other two species of *Spinacia*. These regions therefore probably had an ancestral low recombination rate in both sexes. As suggested above, the recombination pattern must have changed in *S. tetrandra* males to form the older Stratum 1, because (as well as pericentromeric or centromeric regions) this includes a similar sized region that is a rPAR in the other two species (Fig. 2), and presumably in *S. tetrandra* females.

Regions with widely different recombination rates might potentially have different mutation rates. It is thus important to test whether these differences in the likely ancestral recombinational states of the Stratum 2 and 1 regions could affect our strata inferences. To test this, we analysed the Y-X divergence alongside estimates of inter-species divergence. For neutral sites, both divergence values should reflect the mutation rate, and the two *K_S_* values should be correlated, though different genes might differ in their divergence values. To analyse the two sets of divergence values, we used a set of orthologous genes found on the *S. tetrandra* X and Y, and also on the *S. turkestanica* X, and estimated the inter-species divergence between the two X-linked sequences and the Y-X divergence within *S. tetrandra*. A small proportion of the *K_S_* values are very high in both cases. As saturation of the substitutions makes genes with high values uninformative, we analysed divergence sub-sets of the genes, excluding ones with *K_S_*values above different values. With the filtered data, the inter-species values indeed differ between Stratum 1 and 2 genes, and are higher for Stratum 1 than 2, unlike the result without filtering (Table 3, Supplementary Tables 11-13). Also, Stratum 1 still has higher Y-X divergence than Stratum 2, even though removing genes with high Y-X *K_S_* will bias the analysis against detecting strata that evolved before the species split, especially older strata. Overall, therefore, the results support the existence of two strata. Furthermore, the mean *K_S_* values relative to those estimated for inter-species divergence (for the same genes) exceed 1, even after our filtering, which suggests that both strata in *S. tetrandra* evolved before the split of the two *Spinacia* lineages.

**Table 3.** Mean Y-X and inter-species divergence between *S. tetrandra* X-linked alleles and *S. turkestanica* chromosome 4 sequences, estimated as *K_S_* values, and Y-X values corrected using the inter-species divergence values. Results are shown for the entire set of genes satisfying the criterion explained in the text, and for two sub-sets of these genes.

|  | Y-X | Inter-species | Y-X/Inter-species |
| --- | --- | --- | --- |
| **Not filtered (793 genes)** |  |  |  |
| **Stratum 2** | 0.193 | 0.107 | 3.092 |
| **Stratum 1** | 0.298 | 0.101 | 4.660 |
| **Filtered to $K_S < 0.5$ (674 genes)** |  |  |  |
| **Stratum 2** | 0.063 | 0.070 | 1.274 |
| **Stratum 1** | 0.083 | 0.080 | 1.326 |
| **Filtered to $K_S < 0.25$ (634 genes)** |  |  |  |
| **Stratum 2** | 0.050 | 0.052 | 1.214 |
| **Stratum 1** | 0.071 | 0.070 | 1.198 |

Surprisingly, the apparently older stratum 1 is less rearranged between the Y and X than stratum 2. Also, although this Y-linked region is larger than its X counterpart, it has much lower densities of all three types of LTR elements that are abundant in this species’ genome, and has higher gene densities (Fig. 3c, Supplementary Fig. 9). We discuss possible explanations for these findings in the Discussion section. Another unexpected result is that the repetitive sequence density in the *S. tetrandra* SLR is somewhat lower than in the SLRs of the other species (Fig. 2b), even though it has had a longer time for insertions to accumulate, compared with the other two species, as the Y-X divergence, estimated as *K_S_* values in coding sequences, is higher than in *S. oleracea* or *S. turkestanica* (see above). This suggests that sequences have been deleted from Y-linked regions of this chromosome after a period in which repetitive sequences accumulated to high levels. This is consistent with low Y-X synteny and evidence described above for partial genetic degeneration of the *S. tetrandra* Y-linked region, which we next describe in more detail.

To relate the extent of degeneration to the sequence divergence results just described, we followed the approach used for the human XY pair, which infers genes that were on the ancestral chromosome, termed “ancestral genes”) (Wilson Sayres and Makova, 2013). For *S. tetrandra*, we identified genes present in the XLR and also in the corresponding spinach chr4 regions, and counted how many are still present in the homologous *S. tetrandra* YLR, to estimate degeneration levels. We found 1,264 such XLR genes, of which 809 (64%) are still present in the *S. tetrandra* YLR, while about 36% are hemizygous in males (Supplementary Table 14). We estimated gene losses separately for the two evolutionary strata, as one would expect losses to be greater in the older stratum. Of 809 ancestral genes in stratum 1, 606 (74.91%) are still present in the corresponding Y chromosome region, suggesting 25.1% gene losses from this stratum of the Y. In stratum 2, 203 ancestral genes remain on the Y, a lower percentage (44.62%, based on 455 genes), in other words the degeneration level estimated by gene losses is higher in stratum 2 than 1 (Supplementary Table 15).

Degeneration can also be estimated as the proportion of pseudogenes. Among the total of 3,065 genes within the XLR, 181 are pseudogenes (6.0%, see Methods), versus 285 (12.7%) in the YLR (Supplementary Table 16niuibi). The YLR also has a higher proportion of pseudogenes than the XLR among the ancestral genes (see above) that are present on both the *S. tetrandra* XLR and YLR. For stratum 1, 59 of the 606 genes (9.74%) are pseudogenes on the YLR, versus 42 of the 809 XLR genes (5.19%). For stratum 2, with 366 ancestral genes on the X, 41 (9.01%) are pseudogenes, and 59 of the 203 that remain on the YLR (29.06%). Thus both degeneration estimates are higher in stratum 2 than 1 (Supplementary Table 17).

Although all three *Spinacia* species are dioecious, and chromosome 4 is the sex chromosome in all three species, the sex-linked regions (SLRs) are very different (Fig. 4a, b). As outlined in the Introduction, it was previously shown that the SLR of cultivated spinach is a small region (about 12% of chromosome 4) within a very large pericentromeric region with low recombination rates and high repetitive sequence densities in the middle regions of both the X and Y sex chromosomes (She et al., 2023). The situation is similar in *S. turkestanica*. In *S. tetrandra*, however, about 61% of chromosome 4 is completely sex-linked, and there is substantial sequence divergence between the X- and Y-linked alleles of genes across the chromosome, and low coverage in males, indicating genetic degeneration. Much of the chromosome is again pericentromeric, and the X-linked regions are again repeat-rich, just as in *S. oleracea*, but recombination in this region has completely stopped in males, forming a Y-linked haplotype across the most of the pericentromeric region (though not all of it, see **Fig. 2**). Large pericentromeric regions have been detected in many other plants (Brazier and Glémin, 2022) but it has not been clear whether they recombine at low rates, or not at all. Genetic mapping in *S. oleracea* (She et al., 2023) and *Rumex hastatulus* (Rifkin et al., 2021) did not detect recombination, but the rates could be too low to detect without very large family sizes. Low recombination rates can, however, be detected by population genomics approaches. In *S. oleracea*, the absence of diverged Y and X haplotypes across most of the pericentromeric region suggests that enough recombination occurs to prevent sequence differentiation other than accumulation of repetitive sequences, which does not require complete absence of recombination. It is therefore interesting that, in *S. tetrandra*, recombination has stopped in the homologous regions on at least one chromosome arm, changing the region of 4q that we term “pericentromeric PAR” (pPARs in **Fig. 2**) into a fully sex-linked region, and leaving a physically smaller pPAR.

**Figure 4.**
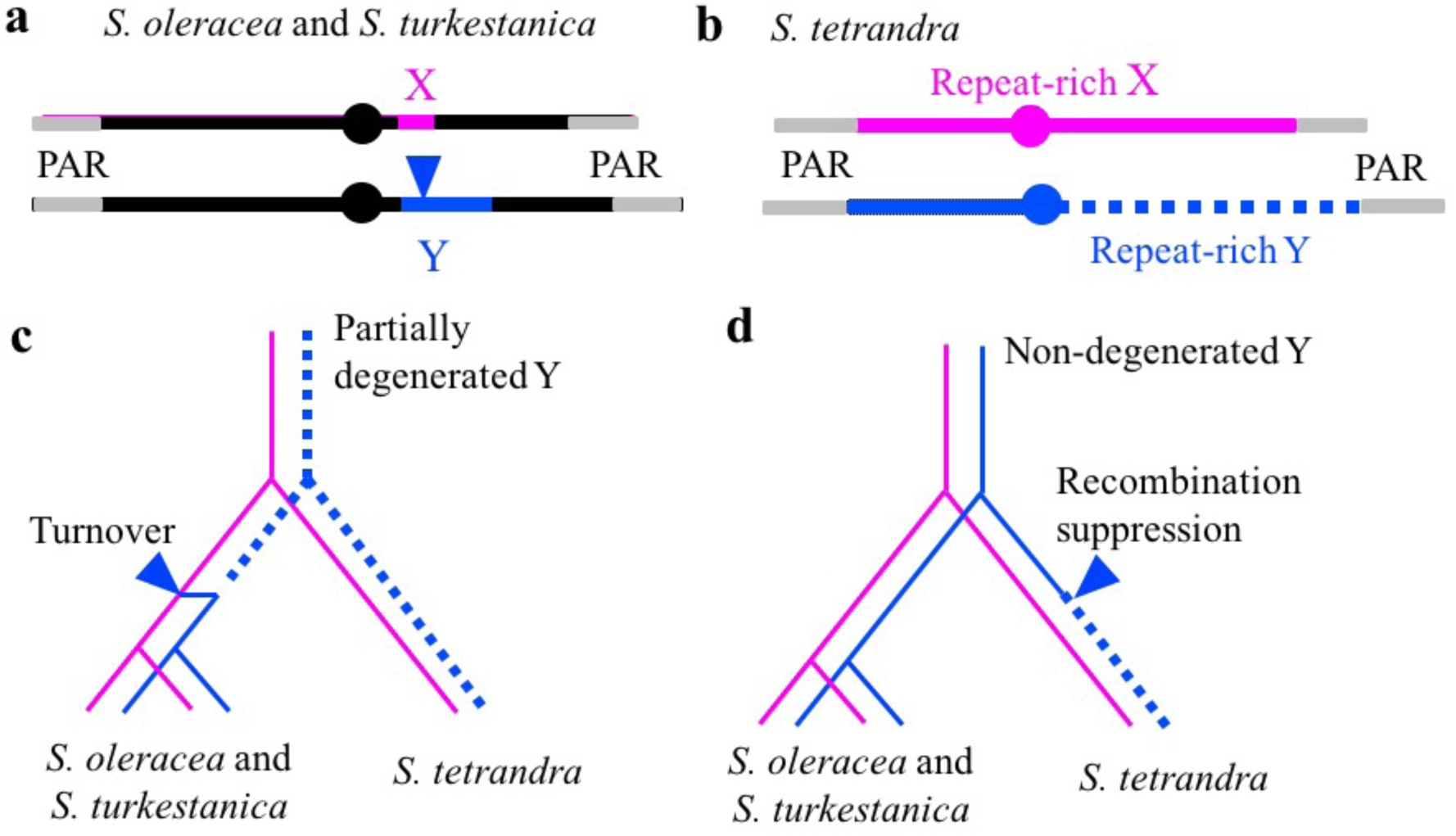
Diagram to illustrate how sex chromosomes could have evolved in the three *Spinacia* species. The structure of sex chromosomes of **(a)** *S. oleracea*/*S. turkestanica* and **(b)** *S. tetrandra*. **(c)** The turnover hypothesis. A new male-determiner arises in a turnover event in an ancestor of *S. oleracea* and *S. turkestanica* whose Y is similar to the *S. tetrandra* Y, in which recombination had stopped, and which is at least partially degenerated (indicated by dotted lines for the Y lineages). The turnover event creates a new Y-linked region that is very similar to the ancestral X state. **(d)** The rapid degeneration hypothesis. The ancestor of all three *Spinacia* species had a small Y-linked region located in an otherwise recombining chromosome, similar to the region in part A. Recombination was suppressed in males across the 4q arm of the sex chromosome in the *S. tetrandra* lineage (two evolutionary strata evolved), and the region degenerated in a time period smaller than the divergence time separating the two.

If the *S. tetrandra* SLR region on the chromosome 4q arm evolved before the split between the two *Spinacia* lineages, our observations are puzzling, at first sight. The Stratum 1 region of high Y-X *K_S_* values has high gene density and high Y-X synteny. In contrast, Stratum 2 to the left of this is rearranged, despite its genes having lower Y-X divergence values. We therefore considered the possibility that the high Y-X *K_S_* values in Stratum 1 are caused by a higher mutation rate than elsewhere on the chromosome, leading to elevated Y-X divergence, and creating a false impression of the age of the stratum. Estimates of inter-species divergence, using *K_S_* values, suggest that the recombinationally active regions of all six chromosomes have higher mutation rates than the pericentromeric regions. The results in Table 3 argue against this possibility, as the inter-species *K_S_* is not much higher in Stratum 1 than 2. This is consistent with the observation that only about half of Stratum 1 evolved from an ancestrally non-pericentromeric region (**Fig. 2**). Overall, the higher Y-X divergence in Stratum 1 genes seems likely to correctly reflect a longer time since recombination stopped, compared with Stratum 2. Both Y-linked regions affected are largely syntenic with their X counterparts. It is not clear whether the inversions found in the older stratum (**Fig. 2**) contributed to its Y-linkage, or whether these occurred after Y-linkage evolved.

To account for *S. tetrandra*’s Stratum 2 region, another recombination suppression event must have subsequently created a new Y-linked region, which has also lost many genes (see Table 4 of gene losses above). This younger stratum spans a large part of the XLR corresponding to the region of the *S. oleracea* chromosome 4q arm extending from the centromere, which was presumably pericentromeric in the ancestor. The lack of synteny could be explained if this region was arranged differently in the haplotype that stopped recombining in male meiosis (and thus became the large younger Y-linked stratum) compared with the arrangement that became the X. The ancestral population cannot be examined to test whether its pericentromeric regions were largely non-recombining and included structural variation, both of which properties are assumed in this scenario.

**Table 4.** Summary of degeneration results, using two different estimates of degeneration.

| Items | Stratum 1 (809 ancestral genes) |  | Stratum 2 (455 ancestral genes) |  |
| --- | --- | --- | --- | --- |
|  | Numbers of genes | Number of pseudogenes among genes present | Numbers of genes | Number of pseudogenes among genes present |
| Number present in YLR | 606 | 59 | 203 | 59 |
| Percentage lost from YLR | 25.1 | 9.74 | 55.4 | 29.06 |
| Number present in XLR | Not applicable (by definition, the X carries all ancestral genes) | 42 | Not applicable (by definition, the X carries all ancestral genes) | 41 |
| Percentage in XLR |  | 5.19 |  | 9.01 |

If recombination became suppressed in both *S. tetrandra* strata before the split from the other *Spinacia* lineage, as argued here, then the sex chromosomes of the other species (**Fig. 4a**) could have evolved from an ancestor that possessed a large SLR similar to that in *S. tetrandra* (**Fig. 4b**). We suggested above that this occurred by a new male-determining factor appearing on the X chromosome (**Fig. 4c**). This scenario is similar to that proposed for a fish, *Poecilia reticulata* (the guppy), whose fully sex-linked region is also physically small, but whose most closely related species, in the genus *Micropoecilia*, have the homologous X chromosome, but highly degenerated Y chromosomes (Charlesworth et al., 2021; Charlesworth et al., 2023; Fong et al., 2023; Metzger et al., 2021).

**Fig. 4d** illustrates an alternative possibility for *Spinacia* (and the guppy), in which the ancestral species had a small YLR on a chromosome that was mostly recombinationally active in males (at least occasionally), and that the species with the degenerated Y chromosome represents a derived state, having evolved large non-recombining regions, or evolutionary strata, which then underwent considerable genetic degeneration since the species split (Darolti et al., 2019; Fong et al., 2023; Metzger et al., 2021). These hypotheses can potentially be tested, because, under the first possibility, the Y-X divergence in the species with the large non-recombining and degenerated region should pre-date the inter-species divergence. As described above, this appears to be the case in *S. tetrandra*, though there are caveats (in the case of the Poecilid fish species, this test could not be used because the extreme degeneration of the YLR left no Y-X pairs for divergence estimation, and the only genes that may be fully Y-linked are closely linked, and physically close, to the PAR (Charlesworth et al., 2021; Fong et al., 2023). As the *S. tetrandra* SLR is less degenerated, it is possible to estimate this divergence value, and test whether it indicates a time for cessation of recombination before the split from the other *Spinacia* lineage, or after the split. As described above, the Y-X divergence (estimated as *K_S_* values) is at least 25% higher than the *K_s_* value estimating the time of the split from either *S. oleracea* or *S. turkestanica*.

## Acknowledgments

This work was supported by the Chinese Academy of Agricultural Sciences Innovation Project (CAAS-ASTIP-IVFCAAS, CAAS-ZDRW202103), and China Agricultural Research System (CARS-23-A-17).

## Competing interests

Authors declare that they have no competing interests.

## Author contributions

W.Q. designed the study. H.S., Z.L., D.C., and F.C., analysed the data. H.S. wrote the manuscript. W.Q., Z.L., H.Z., Z.X. prepared the samples. W.Q., D.C., J.W., X.W., F.C., revised the manuscript. D.C. contributed to interpreting the results, and to writing the manuscript.

## Notes

### Competing Interest Statement

The authors have declared no competing interest.

