## Supplementary Figures for "Two wild *Spinacia* species, *S. turkestanica* and *S. tetrandra*, genomes reveal a sex chromosome turnover in the genus"

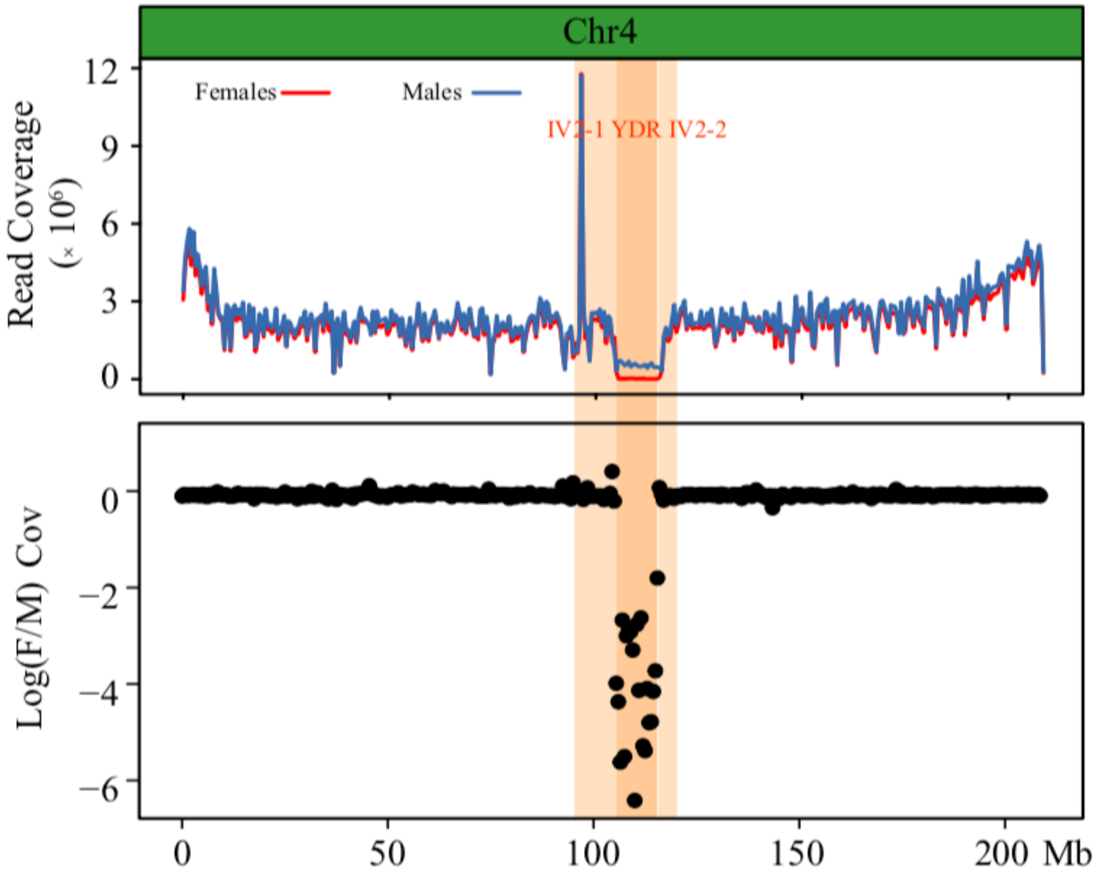


**Supplementary Figure 1. Read coverages of pooled females (red) and males (blue) on the sex chromosome of Tu17S31XY.** Five females and males are used for calculating coverage within each 500-kb window. The yellow background color is putative sex-linked region (SLR) from 95–120 Mb.

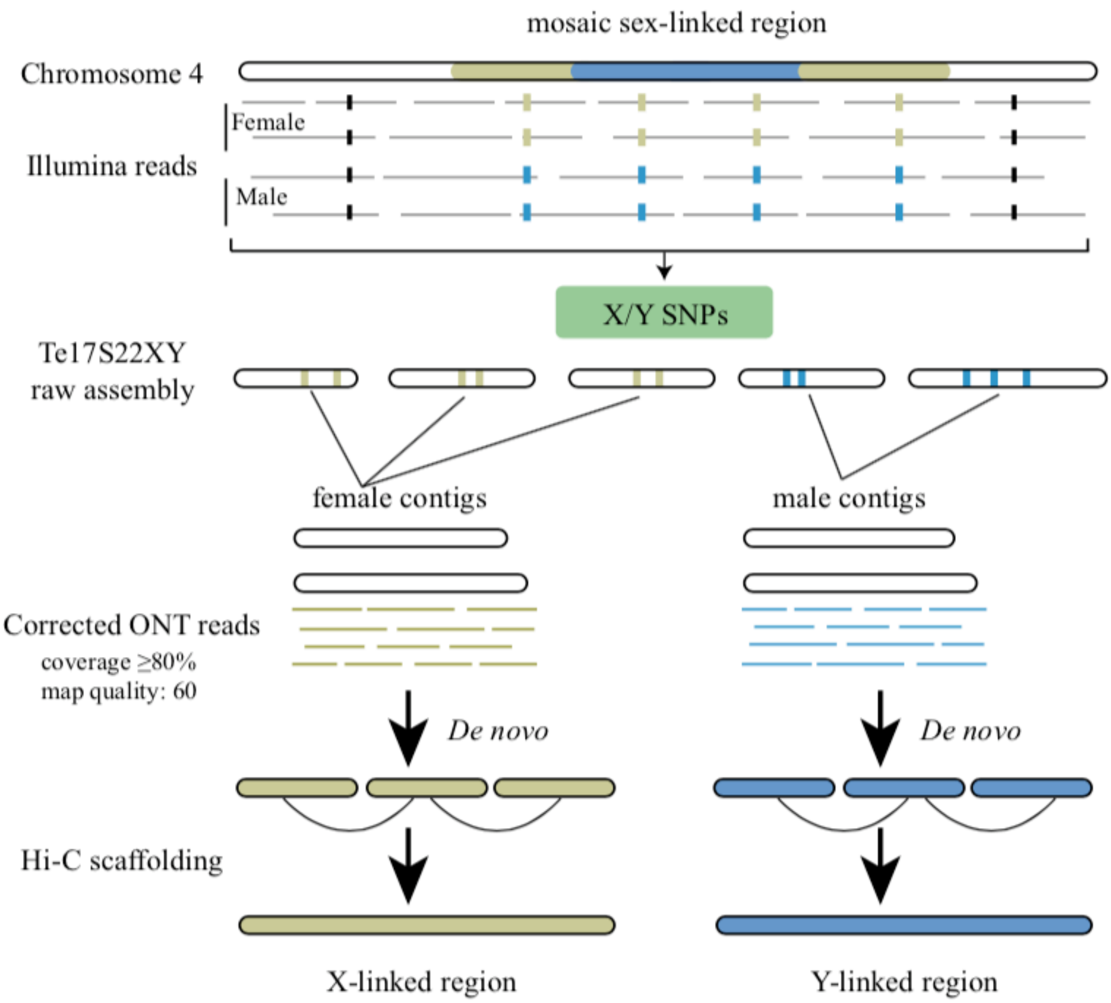


**Supplementary Figure 2. The workflow of X- and Y-linked region assembly in *S. tetrandra*.** Based on the mosaic sex-linked region and X/Y SNPs, female and male contigs from Te17S22XY raw assembly were obtained. Then, female and male ONT reads were generated by mapping corrected ONT reads against female and male contigs using minimap2 with the option '-secondary=no', respectively. Finally, de novo X- and Y-linked contigs using corresponding ONT reads.

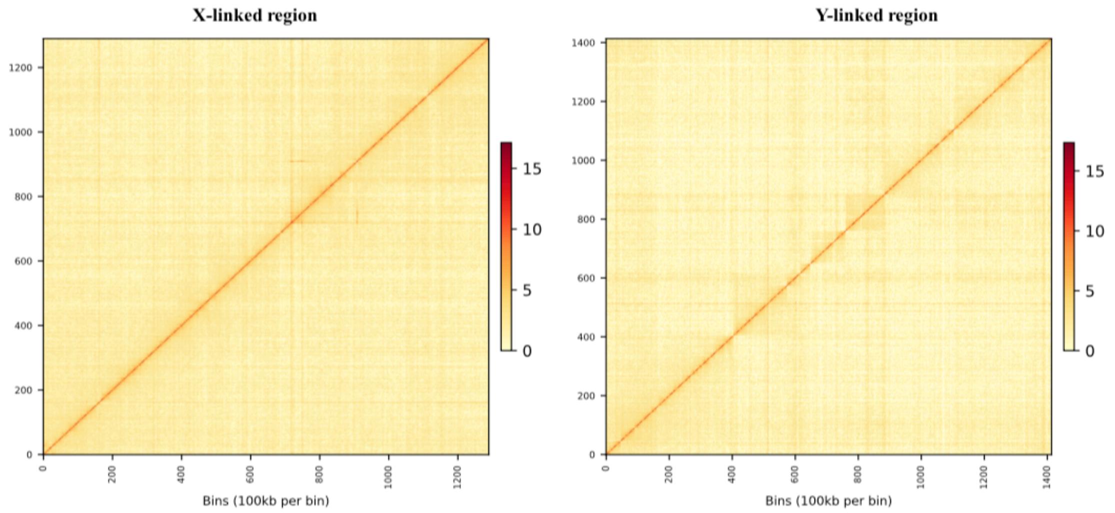


**Supplementary Figure 3. Hi-C contact matrix of the X-linked region and Y-linked region of *S. tetrandra.*** The color intensity represents the frequency of contact between two 100-kb loci.

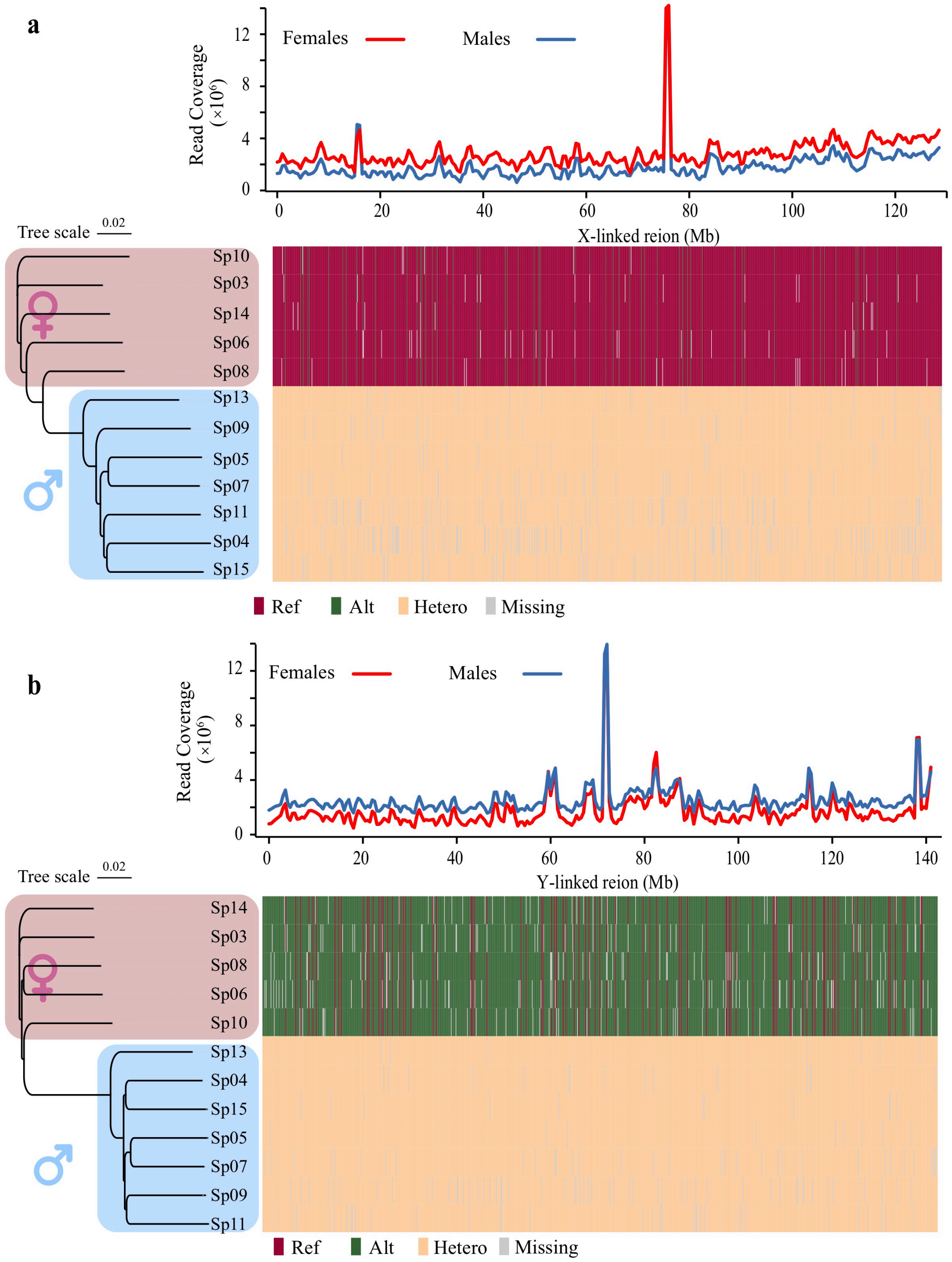


**Supplementary Figure 4. Validation of the large sex-linked region of *S. tetrandra*.** Five females and seven males were used for analyses in 500-kb windows. 19,002 SNPs were homozygous in females, and 45,896 were heterozygous in males, and these SNPs allowed us to phase the sequences into X- and Y-linked haplotypes across the whole of chromosome 4, as illustrated for 1,000 SNPs in the lower figure and in the phylogenetic trees were constructed using these SNPs. Ref: reference; Alt: alternative; Hetero: heterozygous.

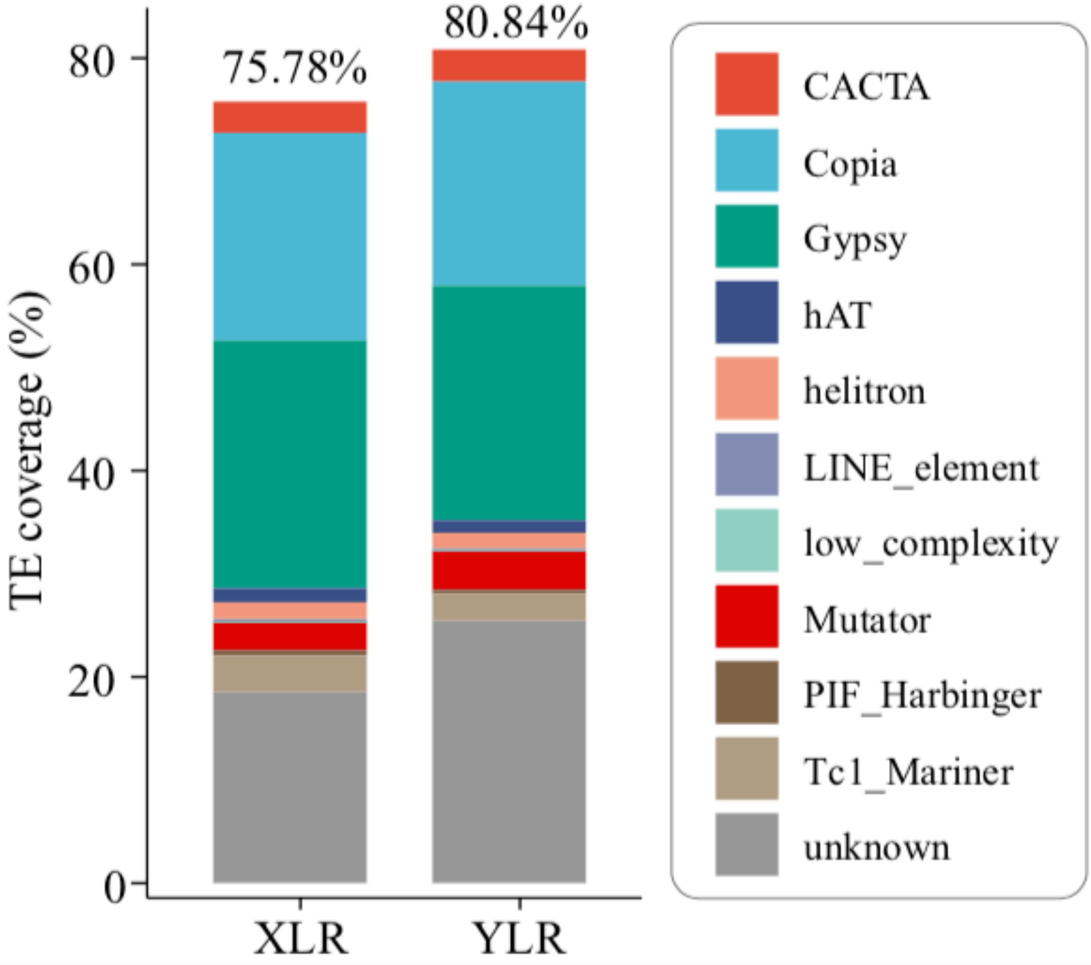


**Supplementary Figure 5. Transposable element (TE) content within the X-linked region (XLR) and Y-linked region (YLR) of *S. tetrandra*.**

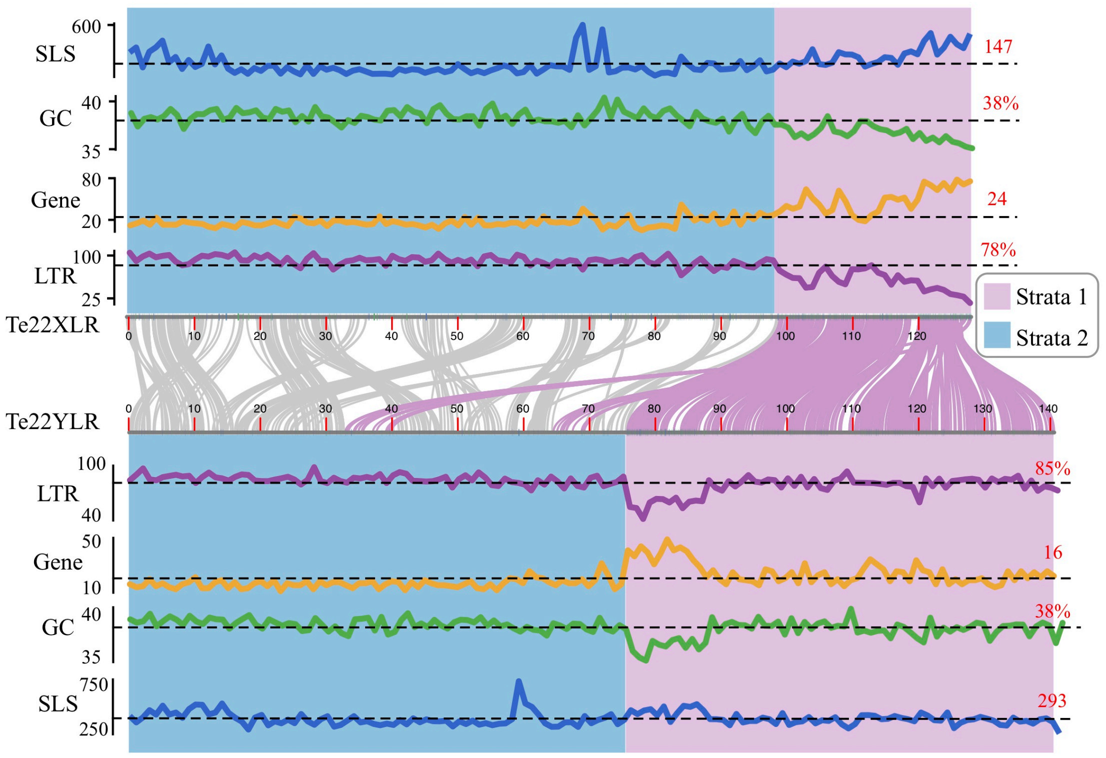


**Supplementary Figure 6. Feature of X-linked (XLR) and Y-linked regions (YLR) of *S. tetrandra*.** Synteny analysis between the XLR and YLR was performed using MCscanX. Long terminal repeat (LTR) and gene density, and GC content, are shown as % values, and the numbers of SNPs showing sex-linkage (SLS) are shown in 1-Mb windows. The dashed lines indicate average values for the X or Y chromosome, and the corresponding values are shown in small red font above the right-hand ends of the lines.

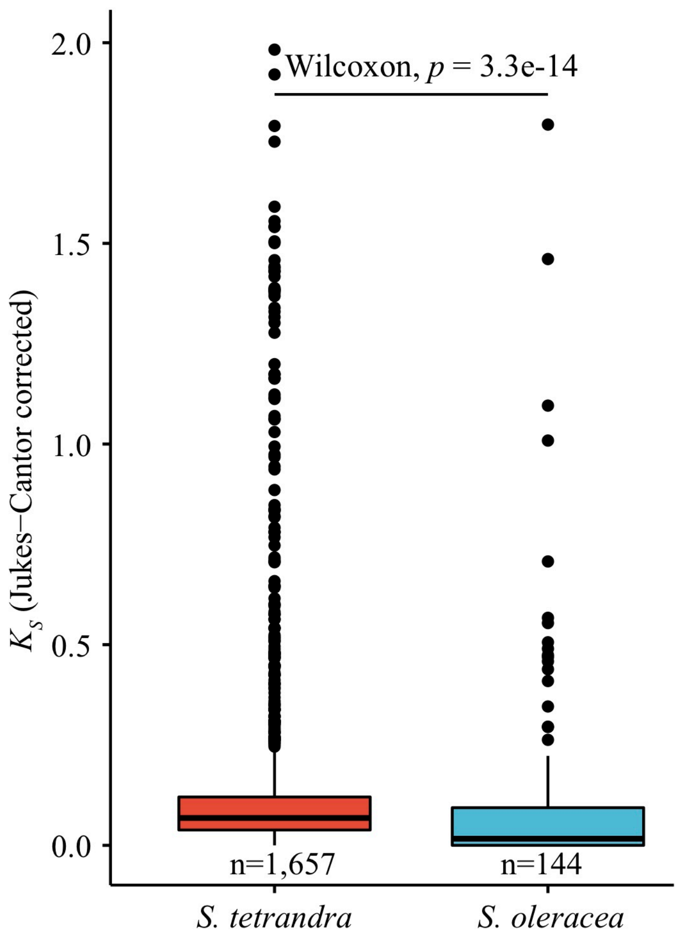


**Supplementary Figure 7. Boxplot of $K_S$ value ($K_S < 2$) of paired XLR/YLR genes in *S. tetrandra* (Te17S22XY) and *S. oleracea* (Sp_YY_v1) assemblies, respectively.** The 144 paired XLR/YLR genes in *S. oleracea* are from previous study (She et al., 2023). In boxplots, lower and upper edges represent 25% and 75% quartiles, respectively, and central lines indicate the median. The whiskers extend to 1.5x the interquartile range. Wilcoxon test ($p$ < 3.3-14) comparing $K_S$ values of paired XLR/YLR genes in the *S. tetrandra* and *S. oleracea* assemblies. XLR: X-linked region; YLR: Y-linked region.

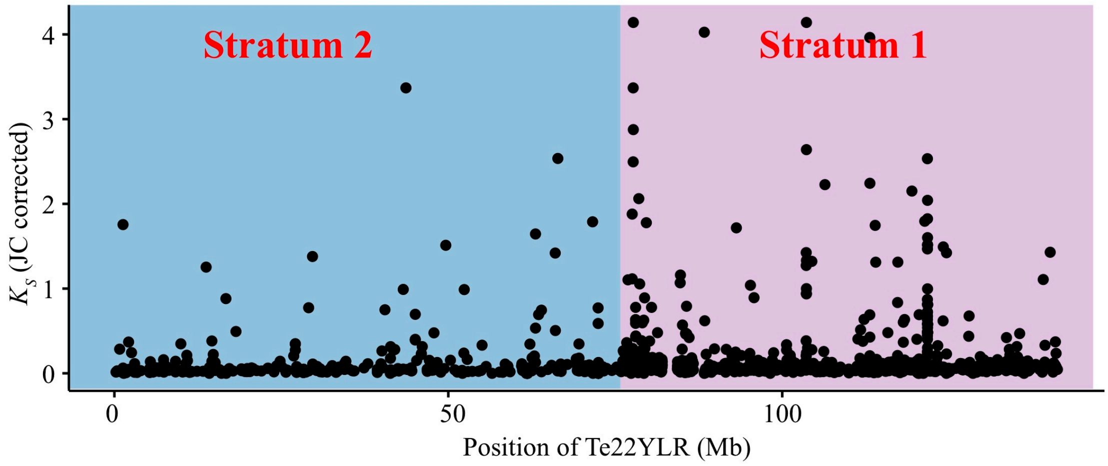


**Supplementary Figure 8. $K_S$ values of allele pairs of genes (no filtered) found in the Te22XLR and Te22YLR sequences.**

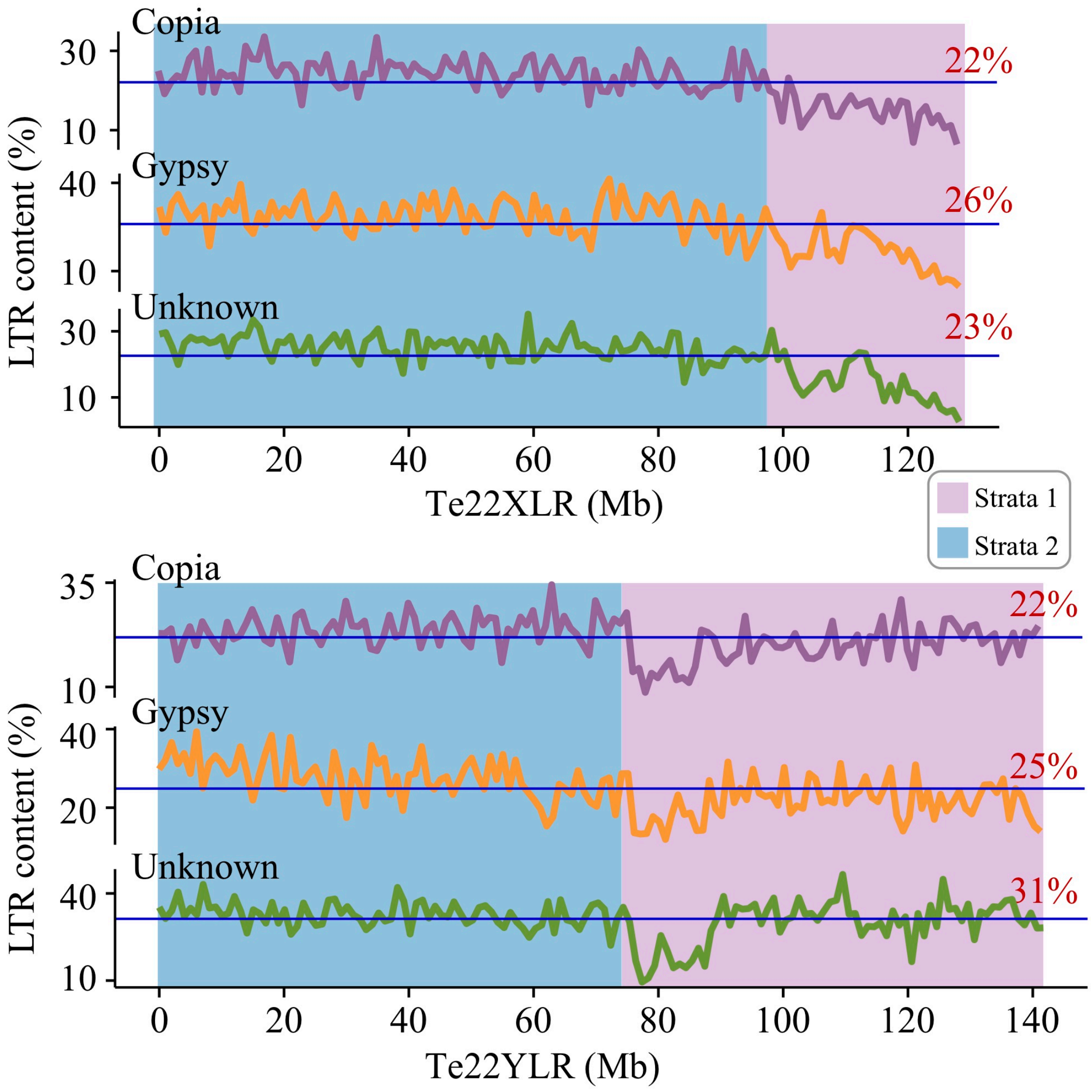


**Supplementary Figure 9. Distribution of three types of LTR content on the XLR and YLR.** LTR content is calculated within the 1-Mb window. The dashed lines indicate X/Y chromosome average and its corresponding value (red) is labeled on the top of the lines.

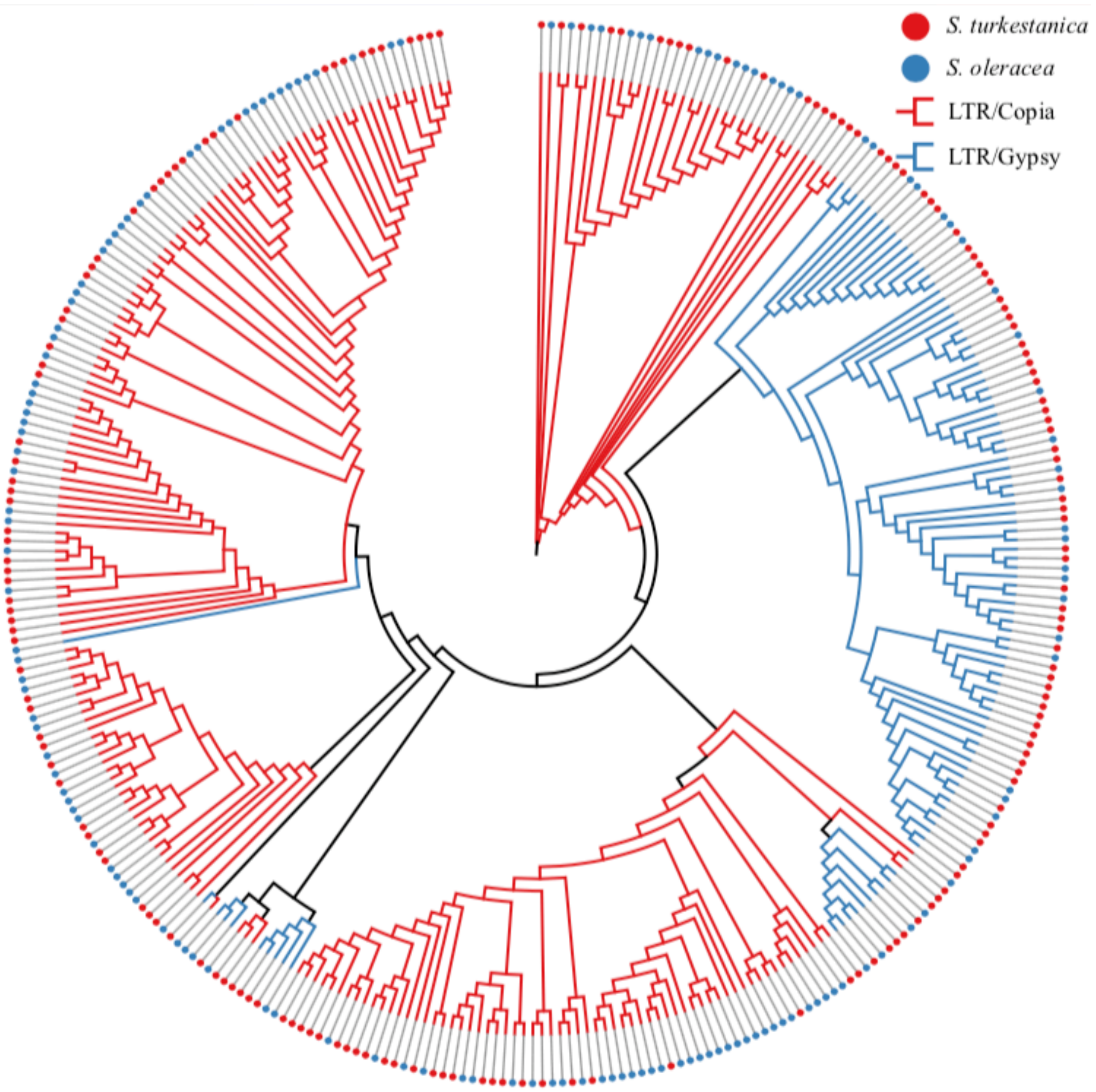


**Supplementary Figure 10. Phylogenetic analysis of *Copia* and *Gypsy* of the YDR in *S. turkestanica* and *S. oleracea*.** We randomly selected 324 LTR orthologs (>90% sequence homology) in the YDR shared by *S. turkestanica* and *S. oleracea* to construct the phylogenetic tree. The LTR sequences are not species-specific, suggesting that the large size of the YDR, with these insertions, evolved before the species diverged.
